# LIN28B-PDZ Binding Kinase Signaling Promotes Neuroblastoma Metastasis

**DOI:** 10.1101/742262

**Authors:** Dongdong Chen, Julie Cox, Jayabhargav Annam, Melanie Weingart, Grace Essien, Komal S. Rathi, Jo Lynne Rokita, Priya Khurana, Selma M. Cuya, Kristopher R. Bosse, Adeiye Pilgrim, Daisy Li, John M. Maris, Robert W. Schnepp

## Abstract

Neuroblastoma is an aggressive pediatric malignancy of the neural crest with suboptimal cure rates and a striking predilection for widespread metastases, underscoring the need to identify novel therapeutic vulnerabilities. We recently identified the RNA binding protein LIN28B as a driver in high-risk neuroblastoma and demonstrated it promotes oncogenic cell proliferation by coordinating a RAN-Aurora kinase A network. Here, we demonstrate that LIN28B influences another key hallmark of cancer, metastatic dissemination. Using a murine xenograft model of neuroblastoma dissemination, we show that LIN28B promotes metastasis. We demonstrate that this is in part due to the effects of LIN28B on self-renewal and migration, providing an understanding of how LIN28B shapes the metastatic phenotype. Our studies reveal that the let-7 family, which LIN28B inhibits, opposes the effects of LIN28B. Next, we identify PDZ Binding Kinase (PBK) as a novel LIN28B target. PBK is a serine/threonine kinase that promotes the proliferation and self-renewal of neural stem cells and serves as an oncogenic driver in multiple aggressive malignancies. We demonstrate that PBK is both a novel direct target of let-7 and that MYCN regulates PBK expression, thus elucidating two oncogenic drivers that converge on PBK. Functionally, PBK promotes self-renewal and migration, phenocopying LIN28B. Taken together, our findings define a role for LIN28B in neuroblastoma metastasis and define the targetable kinase PBK as a potential novel vulnerability in metastatic neuroblastoma.

## INTRODUCTION

Neuroblastoma is an aggressive pediatric tumor of the developing peripheral sympathetic nervous system that remains a substantial challenge in pediatric oncology. (1) At diagnosis, patients with high-risk disease present with striking metastatic burden, underscoring the clinical aggression of this disease. (1) However, our understanding of the molecular mechanisms that drive neuroblastoma metastasis remain incompletely understood, as, consequently, do therapies for metastatic disease.

We (2) and others (3) (4) have demonstrated that LIN28B, an RNA binding protein, is highly expressed in neuroblastoma subsets and that this expression is associated with higher stage neuroblastoma and inferior patient outcome. LIN28B, with its paralog LIN28A, plays pivotal roles in regulating multiple processes that shape normal development, including the cell cycle and apoptosis, self-renewal, glycolysis, and oxidative phosphorylation, among others. (5) (6) These same processes are often subverted in a variety of tumors; accordingly, LIN28B is deregulated in multiple tumor histotypes, including cancers of the colon (7) and ovary (8), Wilms tumor (9), germ cell tumors (10), and hepatocellular carcinomas. (11) Targeted LIN28B overexpression in respective murine tissues leads to the development of neuroblastoma (3), colon cancer (11) (12), and liver cancer (11), credentialing LIN28B as a bona fide oncogene. Mechanistically, LIN28B and LIN28A inhibit the let-7 family of microRNAs and also bind directly to a variety of RNA species, including mRNAs, snoRNAs, and long non-coding RNAs. (5) In humans, there are multiple let-7 family members that have been shown to repress a number of targets implicated in cell proliferation and self-renewal, including expression of *RAS*, *MYC*, and *HMGA2*. (5)

We previously demonstrated that LIN28B promotes neuroblastoma proliferation, in part through regulating the expression of RAN GTPase and Aurora kinase A. (13) (14) While alteration of the cell cycle is a seminal characteristic of cancer, multiple hallmarks comprise the malignant phenotype, including metastatic dissemination. (15) Given the positive association of high *LIN28B* expression with advanced stage disease and poorer outcome (2), along with the fact that LIN28B promotes metastasis in the context of esophageal cancer (16) and colon cancer (7), we investigated whether LIN28B and let-7 act similarly in the context of neuroblastoma metastasis.

## MATERIALS AND METHODS

### Cell culture

Neuroblastoma cell lines were obtained from the Children’s Hospital of Philadelphia (CHOP) neuroblastoma cell line bank and 293T cells from System Biosciences (LV900A-1); all were cultured as previously described. (13) The Emory Genomics Core authenticated cell lines for use in experiments (February 2017). Cell lines were discarded after 20-30 passages. *Mycoplasma* testing was performed every 3-6 months using the *Mycoplasma* test kit (PromoCell, PK-CA91-1024) and on an *ad hoc* basis.

### Plasmid and lentiviral preparation

Lentiviral preparation and transduction were carried out as previously described (13). All shRNA constructs, unless noted otherwise, were obtained from Sigma and are listed in Supplementary Table S1. Dr. David Barrett’s laboratory (Children’s Hospital of Philadelphia) provided the lentiviral GFP/luciferase plasmid used in the neuroblastoma dissemination model (17). Mature let-7i was custom cloned into a lentiviral vector by ViGene Biosciences. The plasmids for the PBK 3’UTR and promoter were purchased from Switchgear Genomics and the Emory Integrated Genomics Core generated the PBK 3’UTR let-7 binding site mutant, with primers described in Supplementary Table S1. The Emory Integrated Genomics Core generated pcDNA3.1-MYCN using the primers in Supplementary Table S1.

### *In vivo* tumor dissemination model

Female *NOD-scidIL2rγ^null^* (*NSG*) mice (6-7 weeks old; The Jackson Laboratory) were housed at the Emory University HSRB Animal Facility in sterile cages in 12-h/12-h light– dark cycles. All experimental procedures were Emory IACUC approved. We infected SKNDZ cells with GFP/luciferase virus and then flow sorted cells to generate the SKNDZ-GFP/luciferase cell line model. We subsequently infected SKNDZ-GFP/luciferase cells with control or shLIN28B lentiviruses, creating stable SKNDZ-GFP-Con, SKNDZ-GFP-shLIN28B-1, and SKNDZ-GFP-shLIN28B-3 models. After acclimatizing, mice were randomized to Con, shLIN28B-1, and shLIN28B-3 groups (n=10 for all). Respective groups received 1 million Con, shLIN28B-1, or shLIN28B-2 cells via tail vein injection. Starting 14 days post injection, bioluminescence imaging was performed twice a week. For imaging, 150 mg/kg luciferin was intraperitoneally injected into mice 10 minutes prior to imaging with IVIS Spectrum Imaging Systems (Perkinelmer). Imaging settings remained the same throughout the study and luminescence intensity was measured using Living Image Software (Perkinelmer).

### Real-Time PCR analysis and Western blotting

RNA was isolated from cells and Real-Time PCR analysis were performed as previously noted. (13) We utilized TaqMan and TaqMan microRNA assays (Life Technologies) to quantitate various transcripts as shown in Supplementary Table S1. Western blotting was carried out as previously detailed, with antibodies as noted in Supplementary Table S1. (13)

### Tumorsphere assays

As previously described (18), single-cell suspensions of cells were detached, filtered through a 100 µm cell strainer, and plated in Tumorsphere medium (DMEM/F12 (Gibco) supplemented with 20ng/ml human recombinant epidermal growth factor (EGF, Corning), 40ng/ml human recombinant basic fibroblast growth factor (bFGF, Corning), 1xB27 (Gibco), 1xN2 (Gibco), 0.1 mM beta-mercaptoethanol (Sigma), 2µg/ml heparin (Stem Cell Technologies), and 1% antibiotic-antimycotic (Gemini). We plated 30,000-40,000 cells per well in 6-well ultra-low attachment plates, and importantly, cells did not aggregate under these conditions. (18) Medium was replenished 3-4 days after plating. Seven to nine days after plating, tumorspheres were dissociated and counted.

### Wound migration assays

Approximately 100,000 cells per well were seeded in 96-well ImageLock plates, yielding 90-100% confluency 24 hours after plating. To inhibit proliferation, cells were treated with 2µg/ml mitomycin C (Sigma) for 1 hour and wounds were generated using the Incucyte wound maker. (19) After rinsing with phosphate-buffered saline 3 times, cells were further cultured in the IncuCyte® Scratch Wound Cell Migration and Invasion System (Essen BioScience) and images were taken every hour during the 24 hour incubation. The relative percentage of wound area occupied by migrated cells was calculated relative to the original wound area using IncuCyte ZOOM analysis software (Essen BioScience).

### 3’UTR and promoter luciferase reporter assays

Assays were carried out with the Lightswitch Luciferase Assay System (Switchgear Genomics) as previously detailed. The effects of control and let-7i on the PBK 3’UTR were normalized to effects on actin (which does not contain let-7 binding sites). (13) PBK promoter luciferase assays were carried out as previously described, with PBK promoter luciferase values normalized to CellTiter-Glo luminescence assay values. (20)

### Neuroblastoma patient datasets

To investigate novel gene-gene correlations and to perform Kaplan-Meier analyses, we utilized the R2: Genomics Analysis and Visualization Platform (http://r2.amc.nl). Individual databases used to investigate correlations and survival analyses are found within the R2: Genomics Analysis and Visualization Platform, with appropriate citations in the text and notation of individual databases in the figure legend. To quantify gene expression at different stages of neuroblastoma, the paired-end reads were aligned with STAR aligner v2.5.2b and expression was quantified in terms of Fragments Per Kilobase of transcript per Million mapped reads (FPKM) with RSEM v1.2.28 using hg38 as the reference genome and GENCODE[^3^] v23 gene annotation. Boxplots were generated using the R package ggpubr and represent *PBK* gene level expression on the y-axis and International Neuroblastoma Staging System (INSS Stage) on the x-axis. ANOVA P-value denotes significance of *PBK* expression differences stratified by INSS stage.

### Chromatin Immunoprecipitation Sequencing (ChIP-Seq)

Kelly MYCN ChIP-Seq was performed and analyzed as previously described (20). For NB-1643 and NGP MYCN and all H3K27Ac ChIP-Seq, cell lines were grown in a 150 mm dish to 60-80% confluency, fixed, and pelleted according to the Active Motif protocol (htts://www.activemotif.com/documents/1848.pdf). Immunoprecipitations were performed using 30 μg of cell line chromatin and 6 μg of N-Myc antibody (Active Motif # 61185) or 4 μg of H3K27Ac antibody (Active Motif #39133). Libraries were prepared by Active Motif and sequenced on a NextSeq 500 to a depth of ∼50 M reads (Jefferson University Genomics Laboratory) and data were analyzed as described previously. (20)

### Statistical analyses

Statistical analyses were performed using GraphPad Prism software version 7. One-way ANOVA with Tukey’s post-hoc test was performed to compare the differences among groups. Unpaired student t-test (two tailed) was used when two groups were compared. Data were presented as mean±standard error. Survival analyses in animal study was performed using the methods of Kaplan and Meier.

## RESULTS

### LIN28B promotes neuroblastoma metastatic dissemination

Patients with high-risk neuroblastoma often present with widespread metastases, but the majority of studies employ models that mimic primary tumor growth; consequently, signaling networks that contribute to neuroblastoma dissemination remain incompletely defined. In aggressive adult malignancies, including esophageal (16) and colon cancer (7), LIN28B promotes metastasis, but whether LIN28B plays similar roles in pediatric tumors is unknown. Given these findings, we hypothesized that LIN28B might similarly promote neuroblastoma dissemination. To address this hypothesis, we employed a tail vein model of neuroblastoma metastasis. (21) Similar models have been utilized to help identify oncogenic pathways that contribute to neuroblastoma metastasis. (22) We first generated control and LIN28B-depleted neuroblastoma cell line models that express luciferase/GFP, allowing *in vivo* imaging.

As shown in Fig. 1A, in comparison to control, LIN28B-depleted cell lines expressed lower levels of LIN28B mRNA and protein. We then injected equal numbers of control and LIN28B-depleted cell lines into the tail vein of *NSG* mice; we employed two independent shRNAs directed against LIN28B (21). There were no significant differences in basal levels of cell death between control and LIN28B-depleted cells, in agreement with previous reports (data not shown). (3) Mice were followed over time via IVIS imaging and those within the control cohort first displayed evidence of metastatic outgrowth at day 14, with significant tumor burden noted in the majority of control mice by day 42. At comparable time points, we observed significantly lower tumor burden in mice bearing LIN28B-depleted tumors, compared to the control cohort (Fig. 1B and Supplementary Fig. S1A). By day 63, all mice in the control cohort had died, with lower tumor burden in cohorts bearing LIN28B-depleted tumors (Fig. 1B). Control mice died by day 56, compared to mice bearing tumors in which LIN28B was depleted (days 91 and 164 for shLIN28B-1 and shLIN28B-3, respectively; Fig. 1C). Following death, subsets of mice underwent a limited necropsy. Almost all mice developed widespread liver and mesenteric metastases, sites often observed within neuroblastoma patients. (1) In addition, we observed some mice with occasional skull/brain and lung metastases, sites sometimes seen in neuroblastoma patients, and associated with worse prognoses (data not shown). (23) To assess whether LIN28B-depleted cell line models maintained lower levels of LIN28B expression, we isolated representative control tumors and LIN28B-depleted tumors that had grown in the liver and performed RT-PCR to quantitate *LIN28B* expression. In comparison to control tumors, LIN28B-depleted tumors maintained knockdown of LIN28B (Supplementary Fig.S1B). Collectively, these studies demonstrate that LIN28B promotes neuroblastoma metastasis in the *in vivo* setting and that shRNA-mediated depletion of LIN28B diminishes tumor dissemination and significantly prolongs survival.

**Figure 1.**
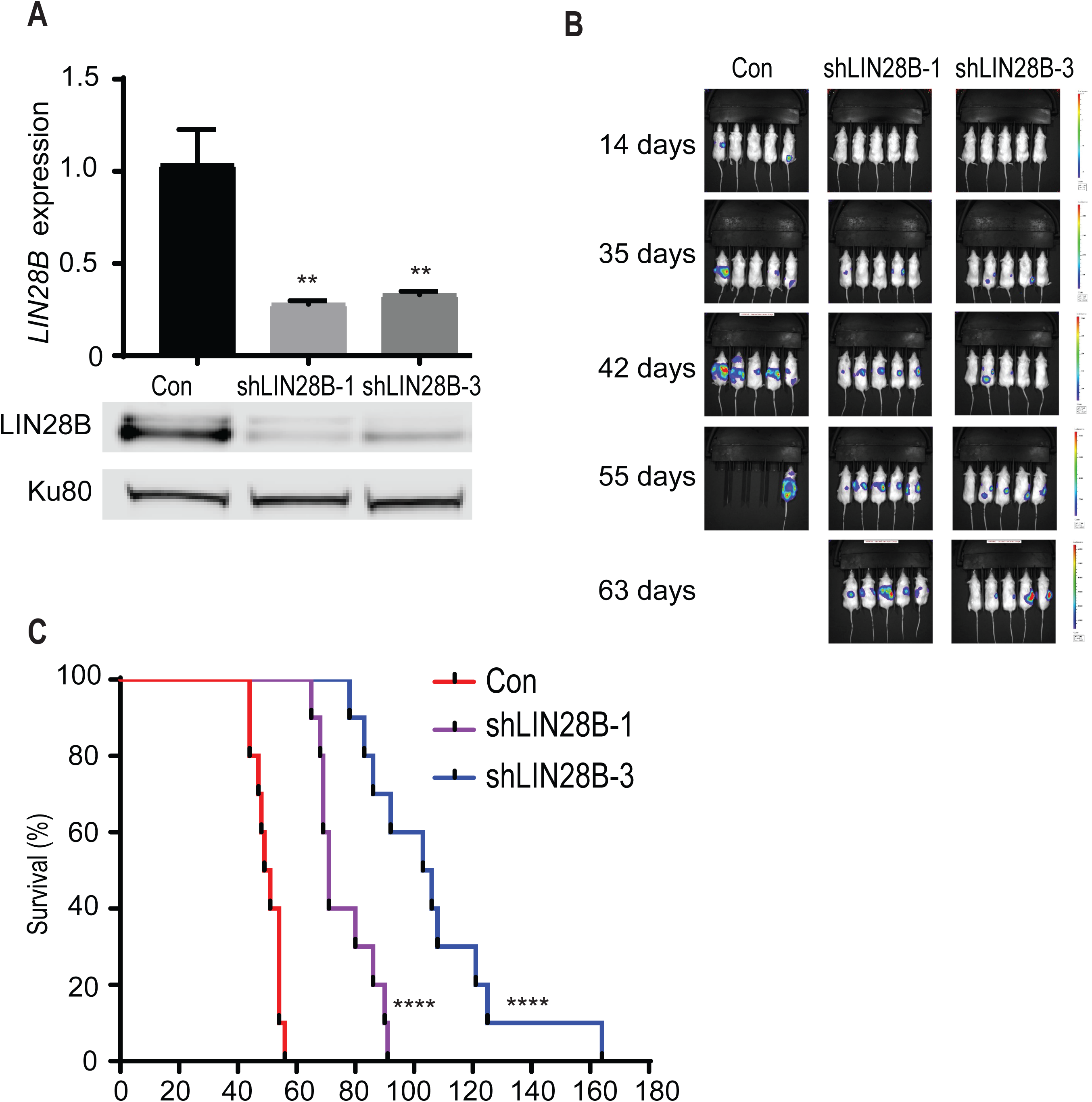
LIN28B promotes neuroblastoma metastatic dissemination. (A) RT-PCR and Western blot demonstrating LIN28B expression in the GFP-luciferase expressing SKNDZ control and LIN28B-depleted cell line models. Two independent shRNAs were used to deplete LIN28B (shLIN28B-1 and shLIN28B-3). Western blotting demonstrates knockdown of LIN28B, with Ku80 as a loading control. (B) IVIS imaging of mice bearing control tumors or tumors in which LIN28B was depleted, at various time points. (C) Kaplan-Meier survival curves for mice bearing control tumors or tumors in which LIN28B was depleted. Error bars represent SEM. ** p<0.01, **** p<0.0001. See also Supplementary Figure S1.

### LIN28B positively influences neuroblastoma self-renewal

Metastasis is a complex process that incorporates multiple steps, including the development of resistance to anoikis, tumor invasion/migration, and metastatic outgrowth. (24) Having demonstrated that LIN28B promotes neuroblastoma metastasis, we next dissected the features of metastasis that LIN28B influences. We first examined the impact of LIN28B on anoikis resistance, but did not observe effects of LIN28B on this property (data not shown). In addition to anoikis resistance, many metastatic cells exhibit enhanced self-renewal, defined as the continuous expansion of cells that possess long-term growth potential (24) (25) For example, N-acetyl-galactosaminyltransferase (GALNT14) promotes breast cancer metastasis by regulating tumor self-renewal. (26) Interestingly, GALNT14 did not influence other properties that traditionally comprise the metastatic phenotype, including anoikis resistance, invasion, or migration. (26) Given the role of LIN28B and let-7 in regulating self-renewal in the context of normal development, we hypothesized that LIN28B might promote self-renewal and thus help drive neuroblastoma metastasis.

We first asked whether LIN28B regulates genes that support self-renewal in both the normal and malignant setting, including NANOG, OCT4, and SOX2, as well as NESTIN, which has been shown to be associated with neural crest stem-like cells as well as with neuroblastoma aggression (27). Neuroblastoma cells were cultured under non-adherent, serum-free conditions and formed tumorspheres. We utilized three independent shRNAs to effectively deplete *LIN28B* (Fig. 2A) and observed decreased levels of *NANOG*, *OCT4*, *SOX2*, and *NESTIN* (Fig. 2B-E). We did not observe a decrease in the levels of *SOX2* for one of the LIN28B-directed shRNAs and speculate that the lower degree of LIN28B knockdown mediated by this shRNA may account for this difference.

**Figure 2.**
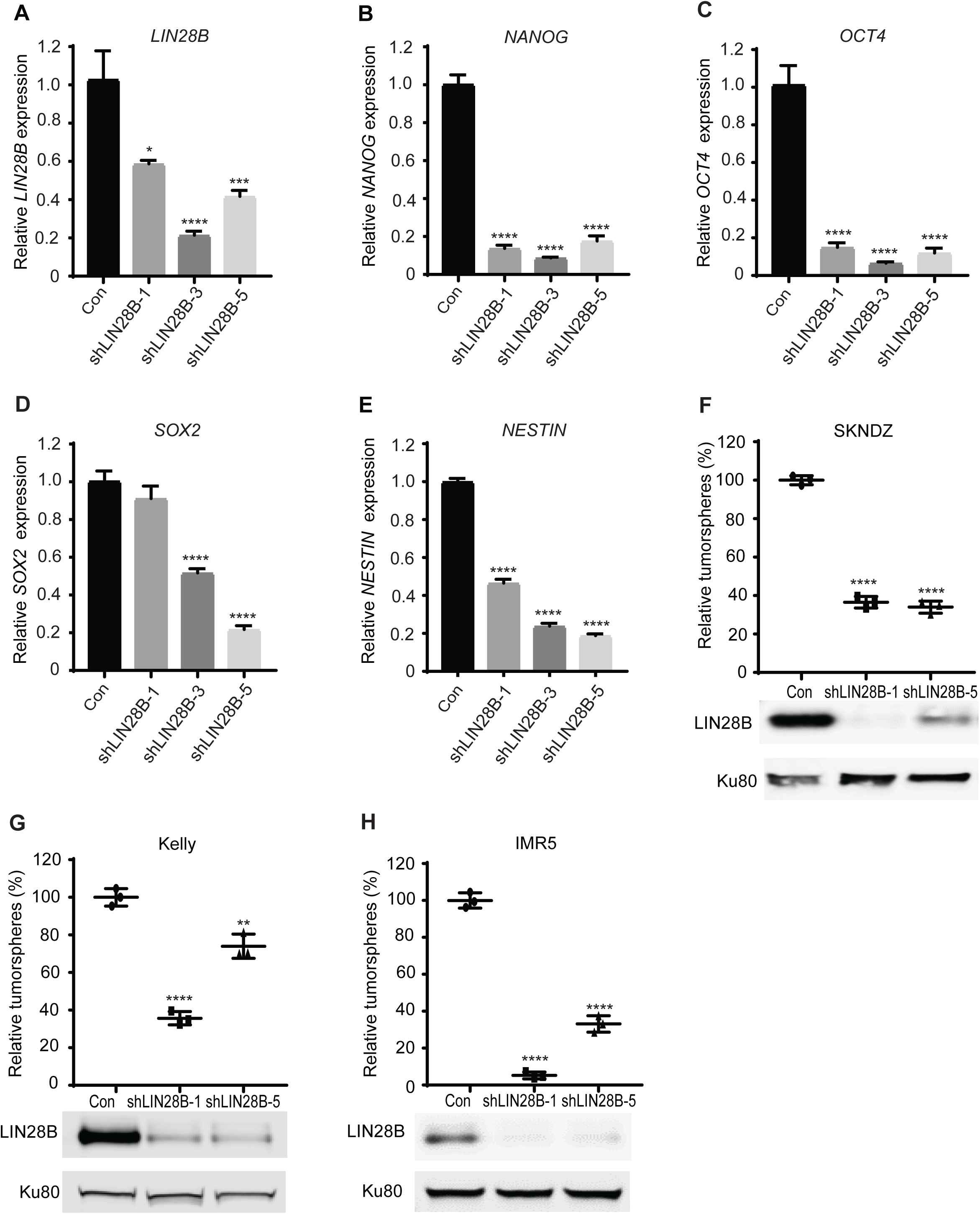
LIN28B positively influences neuroblastoma self-renewal. (A-E) Quantitation of *LIN28B* (A), *NANOG* (B), *OCT4* (C), *SOX2* (D), and *NESTIN* (E), respectively, in SKNDZ cells grown under tumorsphere conditions. (F-H) Tumorsphere quantitation in control and LIN28B-depleted neuroblastoma cell lines SKNDZ (F), Kelly (G), and IMR5 (H). Cell lines were infected with control lentiviruses or lentiviruses expressing two independent shRNAs directed against LIN28B, and grown under tumorsphere conditions. Western blotting of LIN28B and Ku80 in corresponding cell lines. Results are representative of at least two independent experiments. Error bars represent SEM. * p<0.05, ** p<0.01, ***p<0.001, ****, p<0.0001.

The influence of LIN28B on these markers of self-renewal strongly suggests a role for LIN28B in mediating self-renewal. To directly investigate the influence of LIN28B on self-renewal, we employed tumorsphere assays, which provide a functional readout of self-renewal. Previous studies have noted differences between cancer cells grown under adherent, 2-dimensional conditions in comparison to those grown under tumorsphere-forming, 3-dimensional conditions, with some evidence suggesting that tumorsphere cultures may provide a better modeling of the *in vivo* tumor (28) (29). We successfully depleted LIN28B in three neuroblastoma cell lines, as shown by Western blotting and noted decreased self-renewal in all neuroblastoma cell lines (Fig. 2F-H). Collectively, these results suggest that LIN28B promotes self-renewal as one means of promoting metastasis.

### LIN28B promotes the migration of neuroblastoma cells

Another key property that metastatic cells acquire is an enhanced ability to invade and migrate. Given the propensity of high-risk neuroblastoma cells for widespread metastasis (1), we utilized the wound migration assay to determine whether LIN28B contributes to neuroblastoma migration. In two cell line models, we showed that LIN28B indeed promotes neuroblastoma migration (Supplementary Fig. S2A-B). These effects were not due to its influence on cell proliferation, as we noted no differences in cell number between control and LIN28B-depleted neuroblastoma cell lines when assessed at 24 hours (data not shown). However, as a further measure to ensure that the influence of LIN28B on proliferation did not account for effects on cell migration, we treated cells with mitomycin C, which inhibits proliferation. (19) Under these conditions, we again observed that LIN28B depletion impedes neuroblastoma migration (Fig. 3A-C).

**Figure 3.**
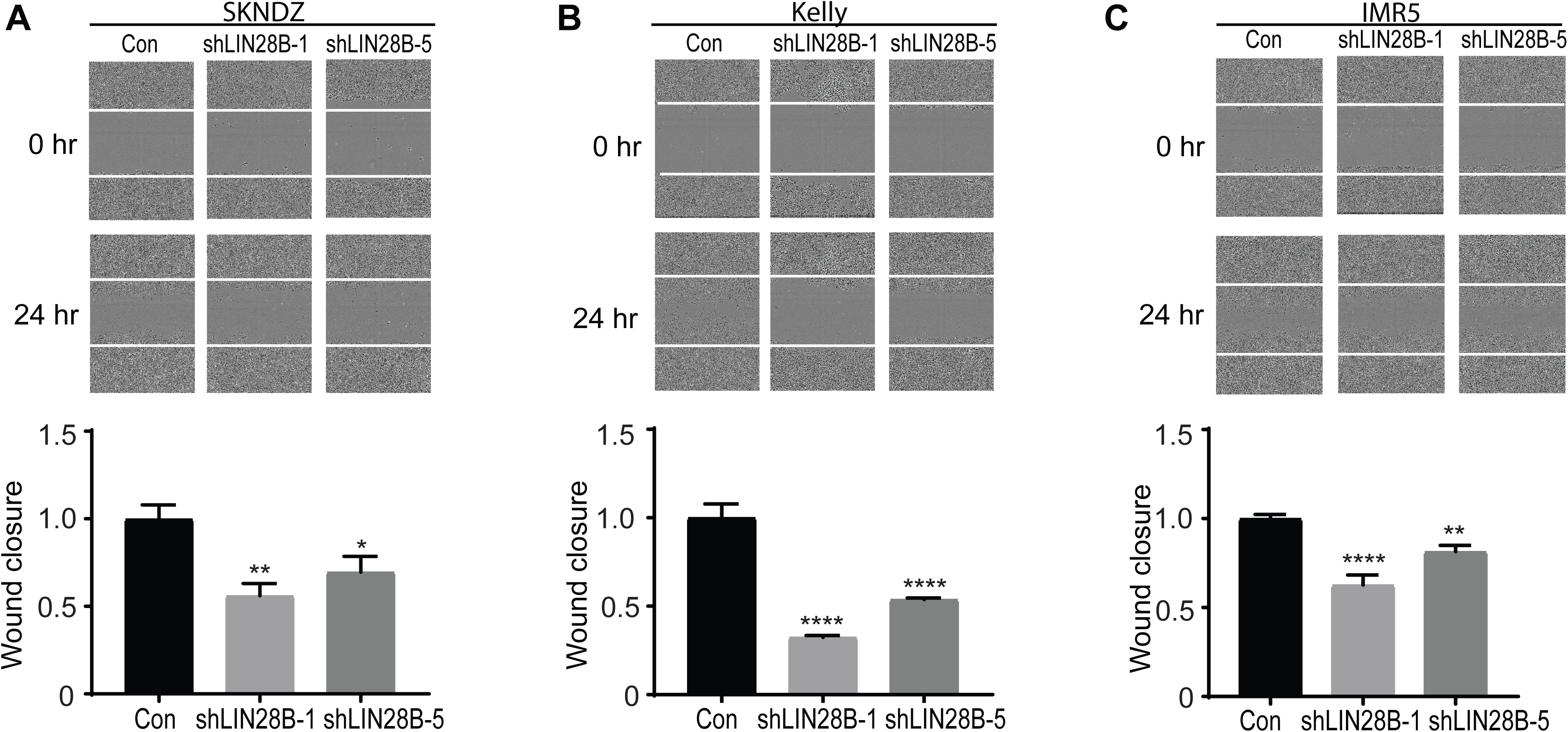
LIN28B promotes the migration of neuroblastoma cells. (A-C) Depicted are representative images from wound closure assays of control and LIN28B-depleted neuroblastoma cell lines SKNDZ (A), Kelly (B), and IMR5 (C). Cell lines were infected with control lentiviruses or lentiviruses expressing two independent shRNAs directed against LIN28B. In addition, given the influence of LIN28B on cell proliferation, cells were treated with mitomycin C. Images were acquired at 0 and 24 hours. Quantitation of wound closure at 24 hours is shown below. Of note, the same cell line models utilized in Figure 2 were used in these assays, with relative LIN28B levels as shown in Figures 2F-H. Results are representative of at least two independent experiments. Error bars represent SEM. * p<0.05, ** p<0.01, ****, p<0.0001. See also Supplementary Figure S2.

### Let-7 inhibits both neuroblastoma self-renewal and migration

One of the most well characterized functions of LIN28B is its inhibition of the maturation and processing of the let-7 family of microRNAs. (5) (6) Therefore, we asked whether let-7 opposes the effects of LIN28B on self-renewal and migration. We generated neuroblastoma cell lines expressing a mature form of let-7i (bypassing the inhibitory effect of LIN28B on let-7 processing) and confirmed overexpression of let-7i (Fig. 4A). Similar to depletion of LIN28B, let-7i overexpression led to significantly decreased self-renewal (Fig. 4B-C). As let-7 inhibits neuroblastoma proliferation (3), we again treated cells with mitomycin and observed that let-7i decreases neuroblastoma migration (Fig. 4D-E). Collectively, these results demonstrate that, with respect to self-renewal and migration, let-7 overexpression mimics the effects seen with LIN28B depletion.

**Figure 4.**
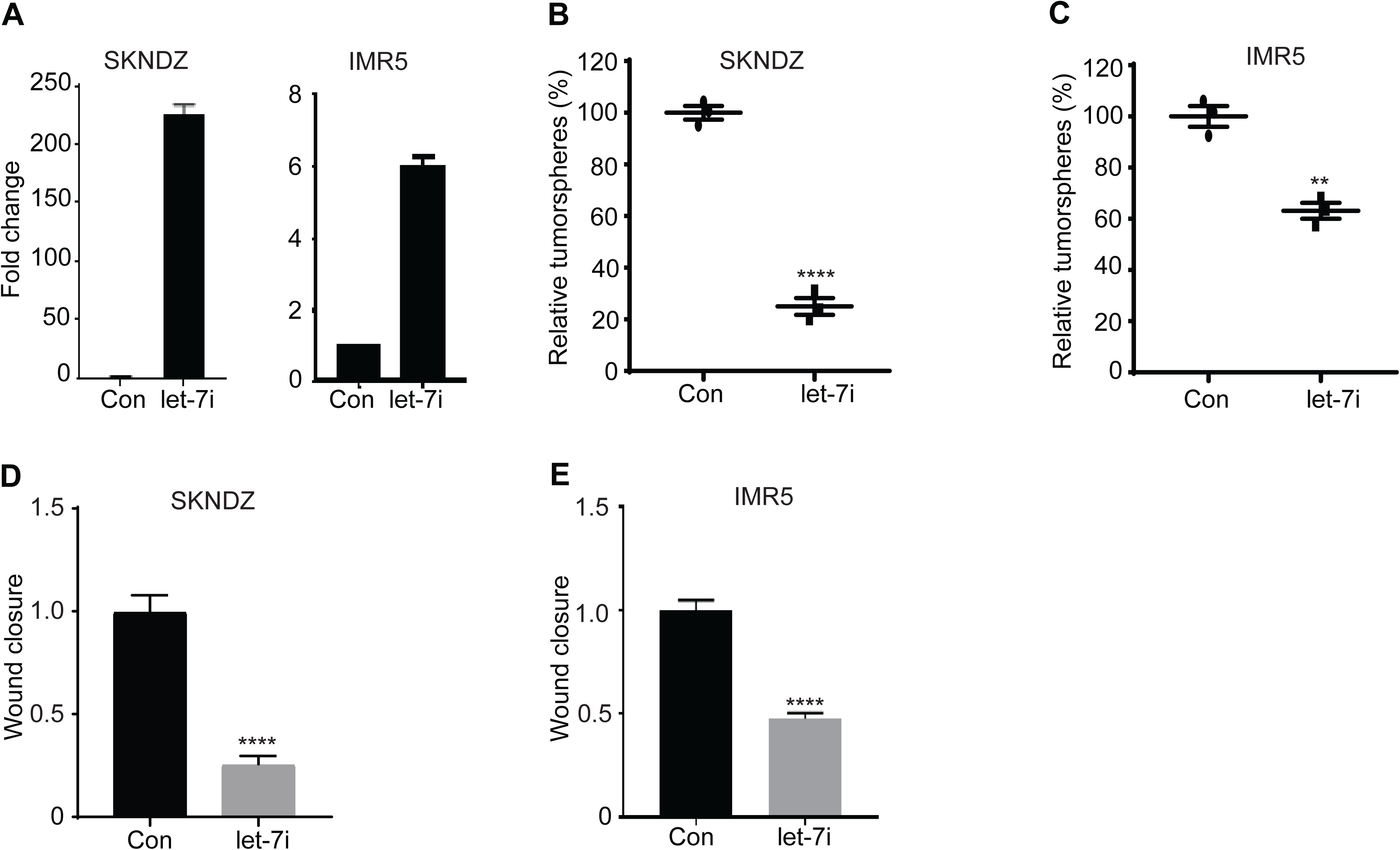
Let-7 inhibits both neuroblastoma self-renewal and migration. (A) RT-PCR demonstrates expression of let-7i in SKNDZ and IMR5 cell lines. Cell lines were infected with control lentiviruses or lentiviruses expressing let-7i. (B-C) Tumorsphere quantitation of control and let-7 expressing neuroblastoma cell lines SKNDZ (B) and IMR5 (C). (D-E) Quantitation of wound closure at 24 hours is shown. Given the influence of let-7i on cell proliferation, cells were treated with mitomycin C. Results are representative of at least two independent experiments. Error bars represent SEM. ** p<0.01, ****, p<0.0001.

### PDZ Binding Kinase is a novel LIN28B-influenced kinase

We next aimed to develop a mechanistic understanding of how LIN28B promotes neuroblastoma dissemination and metastasis. Specifically, we sought to identify novel LIN28B-influenced genes that might be therapeutically targetable. Given our discovery of Aurora kinase A (AURKA) as a novel LIN28B target (13), we hypothesized that LIN28B might coordinate the expression of oncogenic kinases to promote neuroblastoma aggression. (1) Therefore, we first took a candidate approach, perturbing LIN28B levels and investigating effects on kinases that shape the malignant phenotype, including MAPK signaling, MAPK14 (P38) signaling, and PI3K signaling. Interestingly, LIN28B did not influence these oncogenic networks in the context of neuroblastoma (data not shown).

We undertook another approach to identify novel LIN28B-influenced kinases, interrogating the Therapeutically Applicable Research to Generate Effective Treatments project (TARGET; https://ocg.cancer.gov/programs/target) dataset (30), which we had previously utilized in our discovery of RAN and AURKA as novel LIN28B-influenced genes. (13) We evaluated the top 10 kinases significantly and positively correlated with high *LIN28B* expression. We chose to focus our studies on PDZ Binding kinase (PBK), ranked 4/10 of the top correlated kinases, for the following reasons: 1) Its possible role in neuroblastoma, or indeed, in pediatric tumors, was undefined; 2) PBK, also known as T-LAK cell-originated protein kinase (TOPK), is a serine/threonine kinase that promotes the proliferation and self-renewal of neural stem cells (31); 3) PBK is overexpressed in diverse adult histotypes and implicated in multiple hallmarks of cancer, including cell cycle regulation, apoptosis, and metastasis (32); 4) PBK inhibitors have been developed and have demonstrated preclinical efficacy in aggressive adult tumors, including metastatic colon cancer (33) and ovarian cancer. (34) (35)

We found *LIN28B* and *PBK* expression to be highly correlated in neuroblastoma, in both the *MYCN*-amplified (Fig. 5A) and non-*MYCN*-amplified setting (Fig. 5B). (30) To strengthen this observation, we investigated the *LIN28B*-*PBK* correlation in additional neuroblastoma datasets and noted similarly robust correlations (Fig. 5C and Supplementary Fig. S3 A). (36) (37) In further support of a possible oncogenic role for PBK in neuroblastoma, we demonstrated that its expression was associated with higher stage neuroblastoma (Fig. 5D, Supplementary Fig. S3B) (36) (37) and lower overall survival (Fig. 5E, Supplementary Fig. S3C). (36) (37)

**Figure 5.**
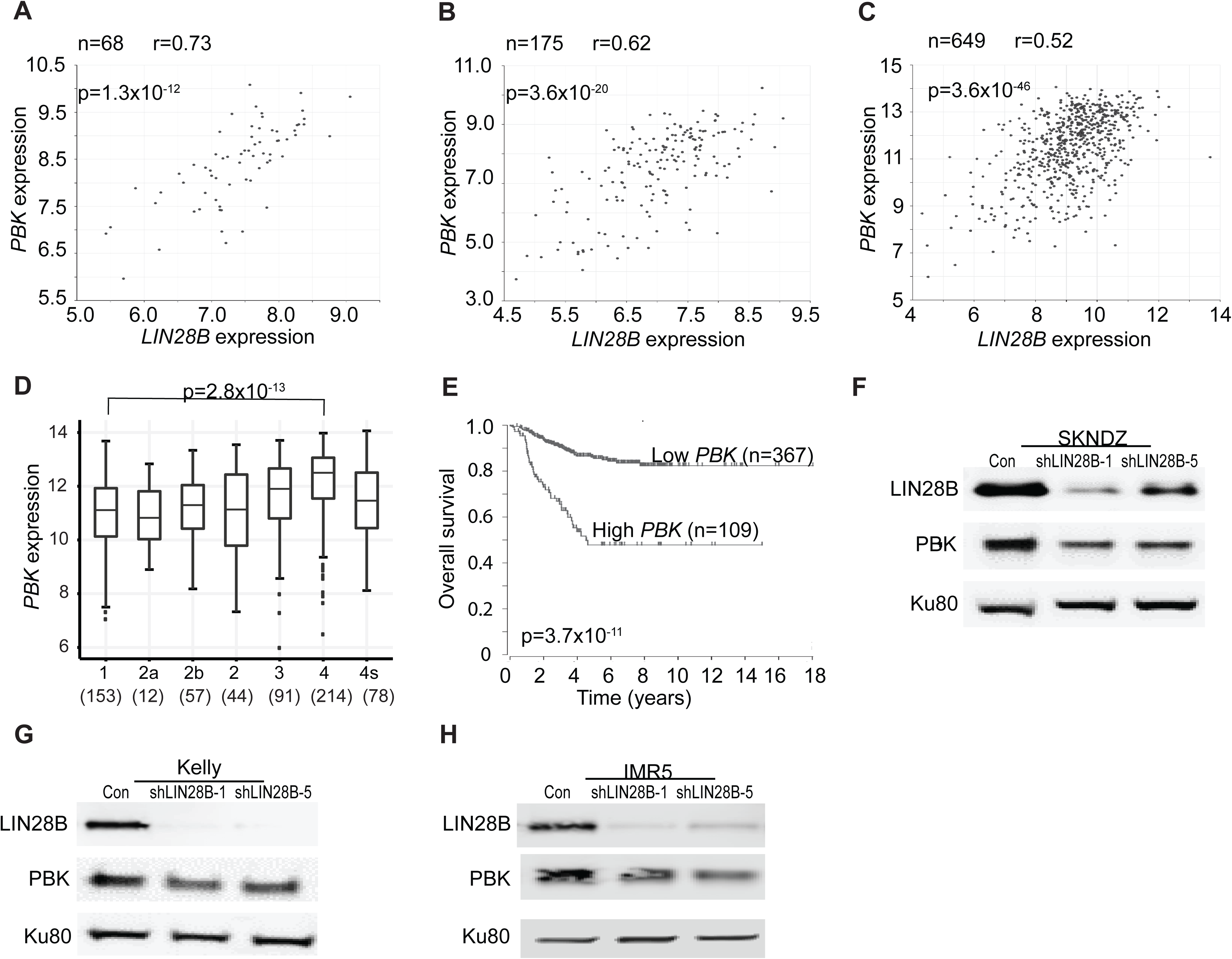
PBK is a novel LIN28B-influenced kinase. (A-B) Correlation between *LIN28B* and *PBK* expression in 68 *MYCN*-amplified (A) and 175 *MYCN* non-amplified (B) primary neuroblastomas. Neuroblastoma datasets obtained from the TARGET consortium. (C) Correlation between *LIN28B* and *PBK* expression in 649 primary neuroblastomas. Data obtained from Kocak dataset. (D) Expression of PBK in primary neuroblastoma tumors, as shown for International Neuroblastoma Staging System stages 1 through 4, with 4S. The number of tumors is depicted in parentheses. Data obtained from Kocak dataset. (E) Kaplan-Meier analysis curves, with individuals grouped by low and high *PBK* expression. Data obtained from Kocak dataset. (F-H) LIN28B and PBK protein levels in SKNDZ (F), Kelly (G), and IMR5 (H) cell lines in which LIN28B was depleted using two independent shRNAs. Ku80 serves as a loading control. P values and r values listed. Error bars represent SEM. See also Figure S3.

While these data demonstrate a robust correlation between *LIN28B* and *PBK* expression, they do not demonstrate whether LIN28B influences the expression of PBK. We depleted LIN28B using two independent shRNAs and noted decreased levels of PBK (Fig 5F-H), demonstrating that LIN28B promotes the expression of PBK.

### Two neuroblastoma oncogenes, LIN28B and MYCN, regulate the expression of PBK

We next dissected the mechanisms by which LIN28B influences PBK expression. As let-7 is a pivotal downstream effector of many LIN28B functions, and as we showed that let-7 inhibits self-renewal and migration, we first investigated whether let-7 influences PBK expression. We engineered three neuroblastoma cell lines to overexpress let-7i and verified significant let-7i overexpression (Fig. 4A and Supplementary Fig. S4A). Western blotting analysis demonstrated that let-7i expression led to downregulation of PBK (Fig. 6A-C). Various microRNA target prediction programs (38, 39) predict that PBK has one let-7 binding site in its 3’UTR and thus we speculated that PBK might be a novel, direct let-7 target. To investigate whether PBK is a novel let-7 target, we performed 3’UTR reporter assays and showed that treatment with let-7i inhibits PBK 3’UTR-driven luciferase activity (Fig. 6D). We then mutated the let-7 binding site in the 3’UTR of PBK and demonstrated that this relieved the inhibitory effect of let-7i, arguing that PBK is indeed a direct let-7 target. Collectively, these experiments demonstrate that one mechanism by which LIN28B regulates the expression of PBK is through let-7.

**Figure 6.**
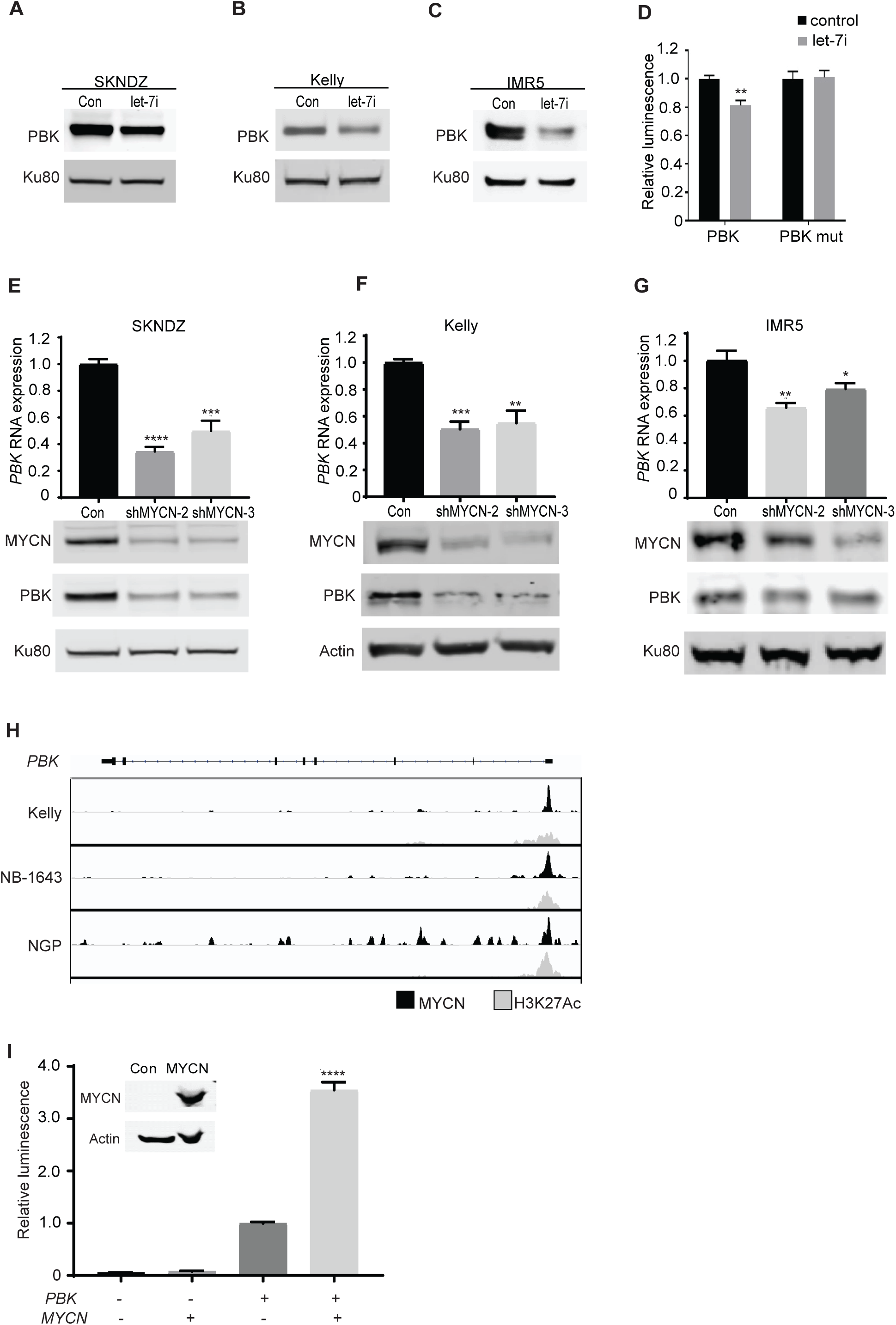
Both LIN28B/let-7 and MYCN directly regulate the expression of PBK. (A-C) PBK protein levels in SKNDZ (A), Kelly (B), and IMR5 (C) cell lines in which let-7i is overexpressed. Ku80 serves as a loading control. (D) 3’UTR assays showing the effect of let-7 on PBK 3’UTR activity. Control microRNA and mature let-7i were transfected into 293T cells and the effect on wild-type PBK 3’UTR and mutated PBK (PBK mut) 3’UTR activity was quantitated. (E-G) Bar graphs depict levels of *PBK* expression in SKNDZ (E), Kelly (F), and IMR5 (G) cell lines in which MYCN was depleted using two independent shRNAs. Immunoblots depict expression of MYCN and PBK protein, with Ku80 serving as a loading control. (H) Plot depicting binding of MYCN (black) and H3K27Ac binding (gray) to *PBK*. ChiP-Seq performed in the *MYCN*-amplified cell line models Kelly, NB-1643, and NGP. (I) PBK promoter assays showing the effect of MYCN on PBK promoter activity. Control and MYCN were transfected into 293T cells and the effect on PBK promoter activity was quantitated. Error bars represent SEM. * p<0.05, ** p<0.01, ***p<0.001, ****, p<0.0001. See also Figure S4.

We and others have previously shown that LIN28B expression is high in *MYCN*-amplified neuroblastoma. (3) (13) LIN28B has been shown to promote MYCN in a let-7 dependent manner (3) and, reciprocally, MYCN binds the LIN28B promoter, positively regulating its expression. (40) Interestingly, PBK is overexpressed in lymphomas and it was recently discovered that MYC positively regulates *PBK* expression by binding to its promoter. (41) We observed that *MYCN* and *PBK* expression are strongly correlated in neuroblastoma tumors (Supplementary Fig. S4B-C). (30) (36) and since MYCN and MYC share many of the same transcriptional targets, we hypothesized that MYCN might directly regulate PBK expression. (42) (43) To determine whether MYCN directly influences PBK, we depleted MYCN using two independent shRNAs, confirming effective knockdown of MYCN at the protein level (Fig. 6E-G). In three independent neuroblastoma cell line models, depletion of MYCN led to decreased PBK levels at both the level of mRNA and protein (Fig. 6E-G). Moreover, analysis of ChIP-Seq data found that MYCN binds to the promoter of PBK in neuroblastoma cell lines, and is accompanied by the active enhancer histone mark, H3K27Ac (Fig. 6H) (20), suggesting direct transcriptional upregulation of PBK by MYCN. To test this hypothesis, we performed PBK promoter reporter assays and showed that treatment with MYCN promotes PBK-driven luciferase activity (Fig. 6I). These results demonstrate that MYCN binds to the promoter of PBK and regulates PBK expression. Taken together, our findings establish that two neuroblastoma oncogenes, LIN28B and MYCN, regulate the expression of PBK and illustrate a convergence of LIN28B/let-7, MYCN, and PBK signaling.

### PBK promotes self-renewal and migration, phenocopying the effects of LIN28B

If one of the major ways by which LIN28B shapes neuroblastoma aggression and metastasis is by positively regulating PBK, then PBK depletion would be expected to phenocopy LIN28B depletion. To assess this hypothesis, we successfully depleted PBK in neuroblastoma cell line models (Fig. 7A-C) and observed that, similar to LIN28B depletion, depletion of PBK led to decreased self-renewal (Fig. 7A-C). Additionally, we found that PBK depletion decreased cell migration, again mimicking the effects of LIN28B depletion (Fig. 7D-E). Thus PBK, a novel LIN28B/let-7 and MYCN target, acts similarly to LIN28B with respect to self-renewal and migration.

**Figure 7.**
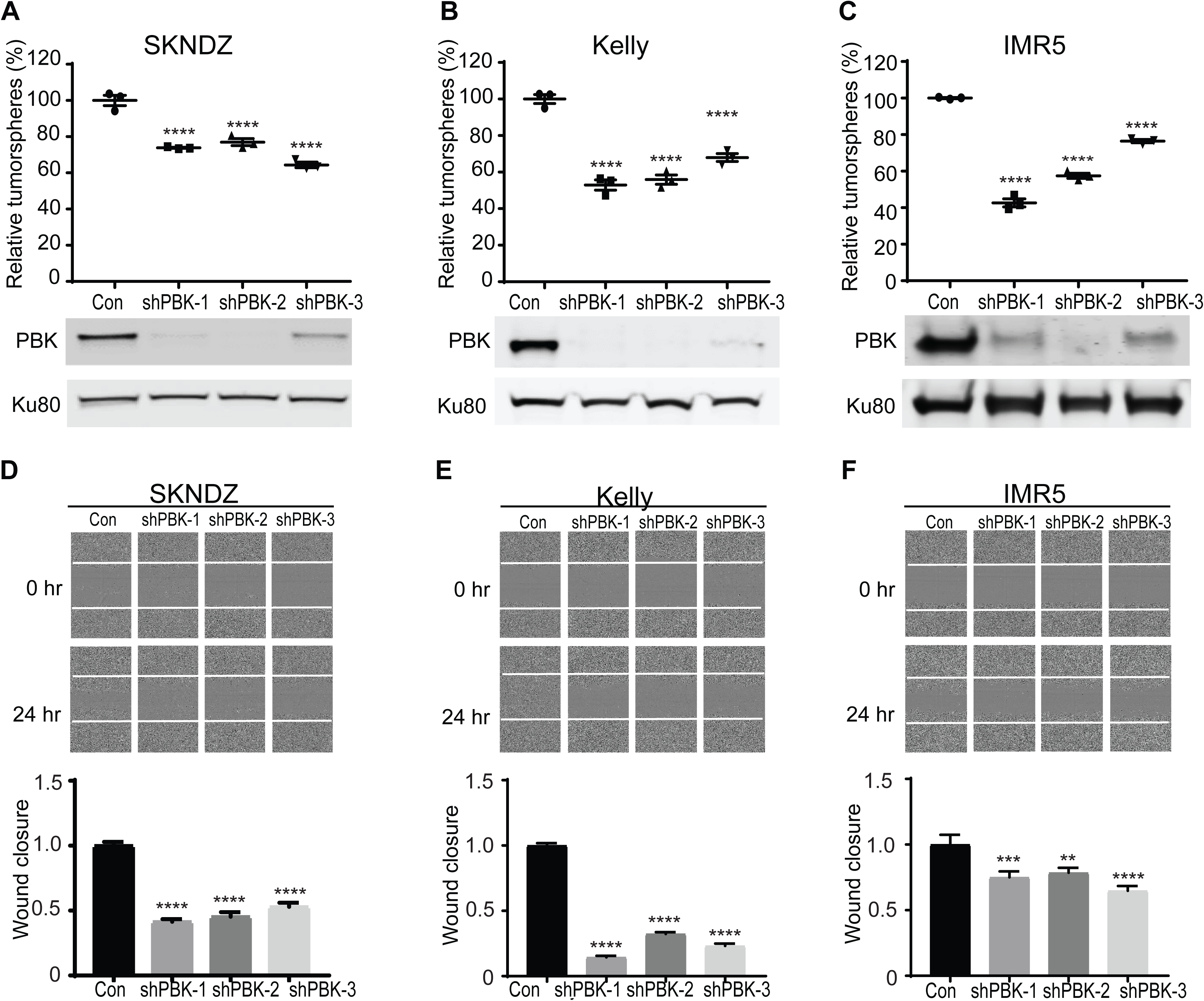
PBK promotes neuroblastoma self-renewal and migration. (A-C) Tumorsphere quantitation in control and PBK-depleted neuroblastoma cell lines SKNDZ (A), Kelly (B), and IMR5 (C). Cell lines were infected with control lentiviruses or lentiviruses expressing three independent shRNAs directed against PBK. Western blotting of PBK and Ku80 in cell lines in corresponding cell lines. (D-F) Depicted are representative images from wound migration assays of control and PBK-depleted neuroblastoma cell lines SKNDZ (D), Kelly (E), and IMR5 (F). Cell lines were infected with control lentiviruses or lentiviruses expressing shRNAs directed against PBK. In addition, cells were treated with mitomycin C. Images were acquired at 0 and 24 hours. Quantitation of wound migration at 24 hours is shown below. Error bars represent SEM. ** p<0.01, ***p<0.001, ****, p<0.0001.

## DISCUSSION

We endeavored to determine how LIN28B shapes neuroblastoma metastasis and to define novel-LIN28B influenced networks that shape aggression. In a neuroblastoma metastasis model, we demonstrate that depletion of LIN28B lessens the dissemination and outgrowth of neuroblastoma. Notably, this model is characterized by widespread liver and mesenteric metastases, along with occasional skull/brain metastases, sites observed in neuroblastoma patients. (1) Our findings expand upon studies demonstrating that targeted expression of LIN28B within the developing murine neural crest leads to the development of neuroblastoma. (3) We show that LIN28B enhances the ability of disseminated neuroblastoma cells to initiate and sustain metastatic colonization and outgrowth. Mechanistically, we demonstrate that LIN28B influences metastasis in part through driving the self-renewal of neuroblastoma cells, potentially allowing tumor cells to repopulate indefinitely. We also show that LIN28B promotes neuroblastoma migration, perhaps allowing cells to exit from the primary tumor and disseminate elsewhere. We demonstrate that let-7 overexpression acts similarly to LIN28B depletion, with respect to self-renewal and migration.

Along with demonstrating the role of LIN28B/let-7 in neuroblastoma metastasis, our studies define PBK as a novel LIN28-influenced kinase. PBK has been shown to be expressed at high levels in multiple aggressive tumors seen primarily in adults, including head and neck cancers, esophageal cancer, liver cancer, colon cancer, and prostate cancer (32). However, its relevance to pediatric tumorigenesis was unknown. Functionally, our studies reveal that PBK promotes neuroblastoma self-renewal and migration, phenocopying the effects of LIN28B. PBK is a tractable therapeutic target, against which clinically relevant inhibitors, primarily targeting the kinase activity of PBK, exist. In preclinical models of colon (33) and ovarian cancer (33), PBK inhibition led to decreased metastatic dissemination. Future studies will focus on elucidating the role of PBK in additional models of neuroblastoma as well as on metastasis within the *in vivo* setting. More broadly, it will be of interest to characterize the expression of PBK in other pediatric malignancies and investigate whether PBK promotes tumor aggression in these contexts.

Mechanistically, our data indicate that LIN28B and MYCN signaling intertwine and influence PBK expression by two different mechanisms. First, our studies reveal PBK to be a novel and direct let-7 target. Second, our data demonstrate that MYCN binds the *PBK* promoter and promotes MYCN expression. This is reminiscent of Aurora kinase A, which we previously identified as a LIN28B/let-7 target and other investigators have shown interacts with MYCN to stabilize MYCN protein. (44) While drugging LIN28B/let-7 is still in its infancy, investigators have developed small molecule inhibitors that disrupt the repression of let-7 by LIN28B, allowing restoration of let-7 levels. (45) Aurora kinase A inhibitors, such as alisertib, have undergone Phase 2 testing in neuroblastoma and were found to be well tolerated and to demonstrate activity, primarily in the non-*MYCN*-amplified context. (46) Additionally, BET inhibition provides a means of targeting MYC/MYCN and has demonstrated efficacy in preclinical neuroblastoma models. (47) Moreover, others have shown that MYCN influences and collaborates with epigenetic machinery, including the PRC2 complex, providing additional therapeutic opportunities. (50) Finally, PBK inhibition has demonstrated *in vivo* efficacy against aggressive adult histotypes. (32) Due to the inherent heterogeneity within primary tumor/metastatic sites, as well as the heterogeneity shaped by multimodal neuroblastoma therapies, developing a robust compendium of therapeutic agents for combinatorial regimens will likely be necessary to optimize neuroblastoma therapy. It will be of substantial interest to determine whether regimens targeting LIN28B/let-7, PBK, MYCN, and/or AURKA might improve the treatment of neuroblastoma aggression and metastasis.

## AUTHOR’S CONTRIBUTIONS

**Conception and design:** D. Chen, J. Cox, R.W. Schnepp

**Development of methodology:** D. Chen, J. Cox, J. Annam, R.W. Schnepp

**Acquisition of data:** D. Chen, J. Cox, J. Annam, M. Weingart, G. Essien, P. Khurana, S.M. Cuya, A. Pilgrim, D. Li, R.W. Schnepp

**Analysis and interpretation of data (biostatistics, statistical analysis, interpretation of clinical data and genomic datasets):** D. Chen, K.S. Rathi, J. L. Rokita, K. Bosse, A. Pilgrim, R.W. Schnepp

**Writing, review and/or revision of the manuscript:** D. Chen, R.W. Schnepp

**Administrative, technical, or material support:** J.M. Maris, R.W. Schnepp

**Study supervision:** R.W. Schnepp

## ACKNOWLEDGMENTS

This work was supported in part by NIH Grant K08-7K08CA194162-02 (R.W.S.), NIH Grant R35 CA220500 (J.M.M.), NIH Grant R01 CA124709 (J.M.M.), the Alex’s Lemonade Stand Foundation (J.L.R., K.R.B., R.W.S.), the Damon Runyon Cancer Research Foundation (PST-07-16; K.R.B.), Hyundai Hope on Wheels (R.W.S.), Andrew McDonough B+ Foundation (R.W.S.), Team Connor Foundation (R.W.S.), Rally Foundation for Childhood Cancer Research (R.W.S.), the Winship Cancer Institute American Cancer Society Institutional Research Grant (R.W.S.), the Aflac Cancer and Blood Disorders Center Trust (R.W.S.), and the William Woods, M.D., Aflac Clinical Investigator Chair (R.W.S.).

In addition, we thank Dr. David Barrett, the Children’s Hospital of Philadelphia, for the lentiviral GFP/Luciferase plasmid. Finally, we thank Dr. Oskar Laur and the Emory Custom Cloning Core/Emory Integrated Genomics Core for the cloning of various constructs. The Emory Integrated Genomics Core is subsidized by the Emory University School of Medicine and is one of the Emory Integrated Core Facilities. Additional support was provided by the National Center for Advancing Translational Sciences of the National Institutes of Health under Award Number UL1TR000454. The content is solely the responsibility of the authors and does not necessarily reflect the official views of the National Institutes of Health.

## SUPPLEMENTARY FIGURE LEGENDS

**Supplementary Figure S1, related to Figure 1.**
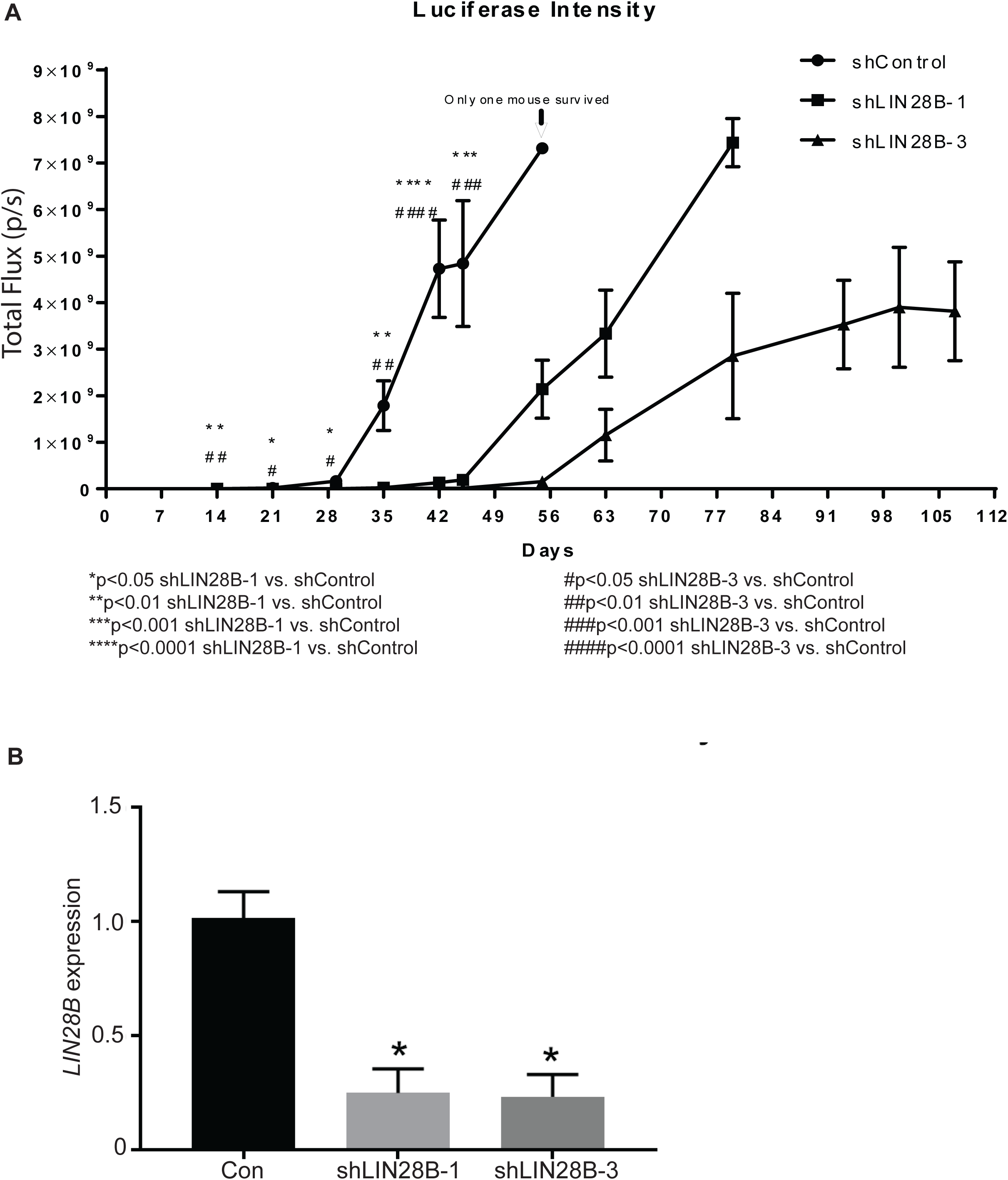
LIN28B promotes neuroblastoma dissemination in the *in vivo* setting. (A) Mean bioluminescence intensities of mice from control and shLIN28B-depleted cohorts. (B) RT-PCR demonstrating LIN28B expression in the tumor tissue of control mice or mice with tumors in which LIN28B was depleted. Of note, we isolated 3 control tumors, 2 tumors from the shLIN28B-1 cohort, and 2 tumors from the shLIN28B-3 cohort. * p<0.05.

**Supplementary Figure S2, related to Figure 3.**
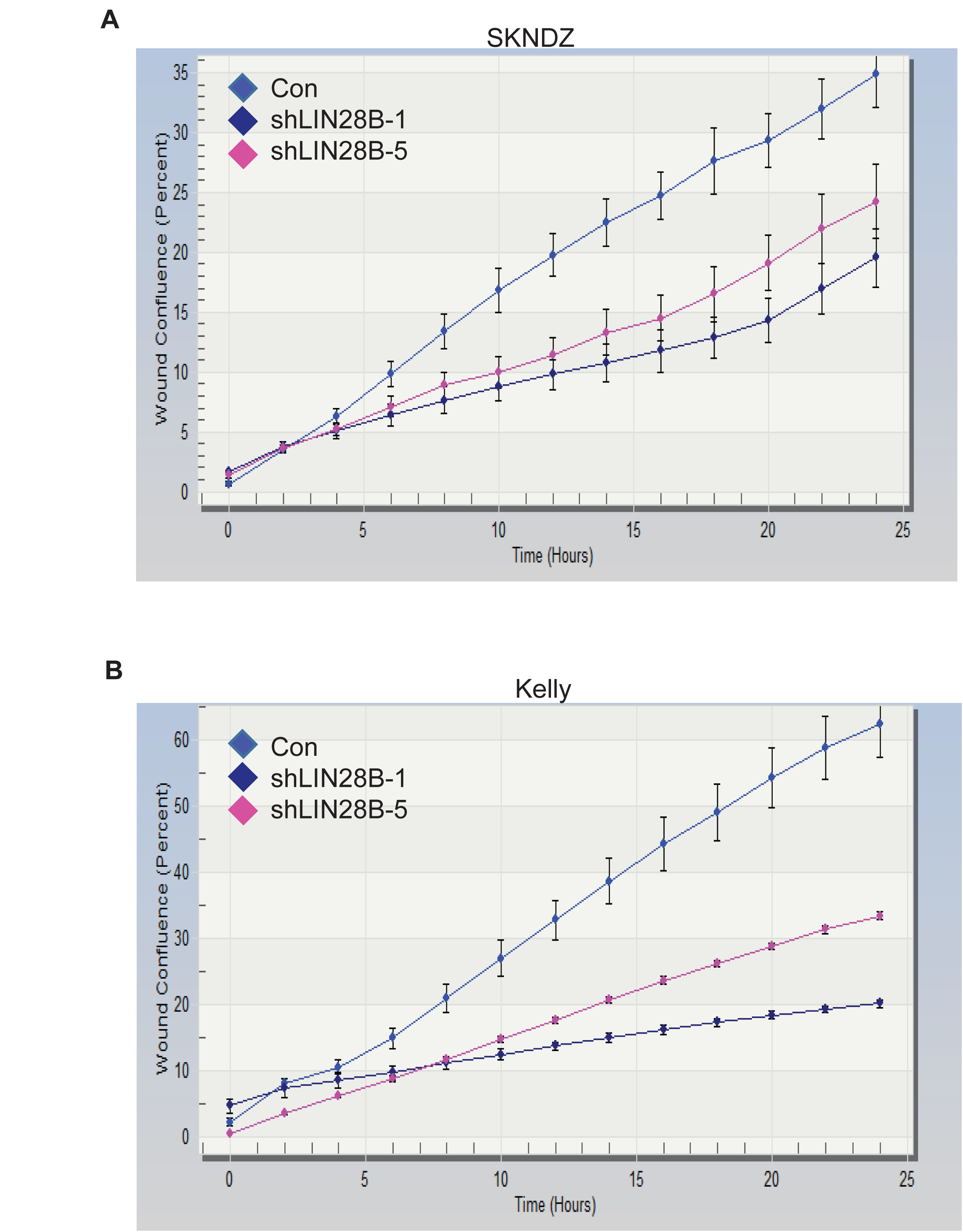
LIN28B promotes the migration of neuroblastoma cells. (A-B) Quantitation of wound closure over 24 hours for SKNDZ (A) and Kelly (B) cells infected with control lentiviruses or lentiviruses expressing shRNAs directed against LIN28B. Of note, cells were not treated with mitomycin. Error bars represent SEM.

**Supplementary Figure S3, related to Figure 5.**
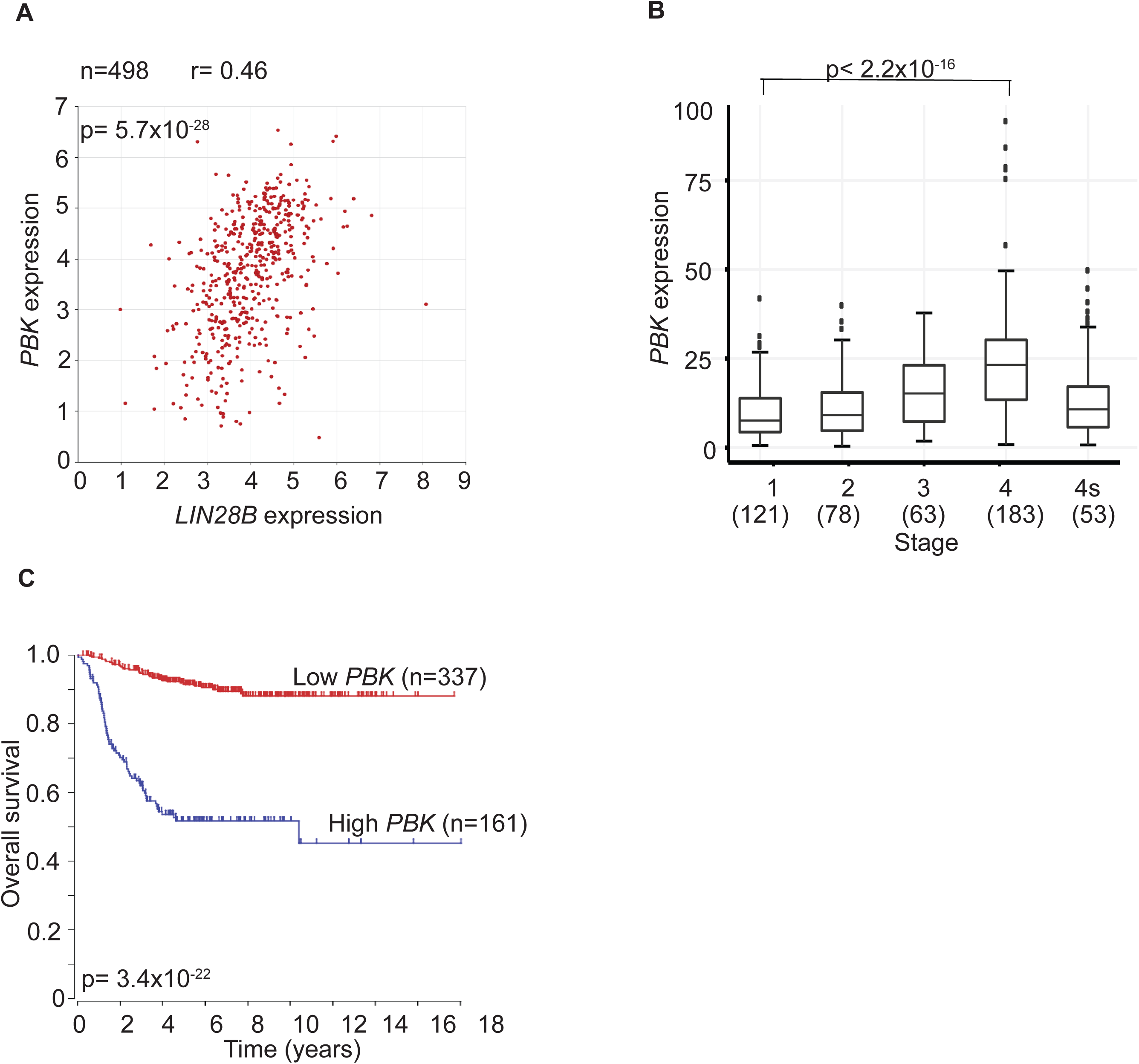
PBK is a novel LIN28B-influenced kinase. (A) Correlation between *LIN28B* and *PBK* expression in 498 primary neuroblastomas. (B) Expression of *PBK* in primary neuroblastoma tumors, as shown for International Neuroblastoma Staging System stages 1 through 4, with 4S. The number of tumors is depicted in parentheses. (C) Kaplan-Meier analysis curves, with individuals grouped by low and high *PBK* expression. Data obtained from the Su dataset. P values and r values listed.

**Supplementary Figure S4, related to Figure 6.**
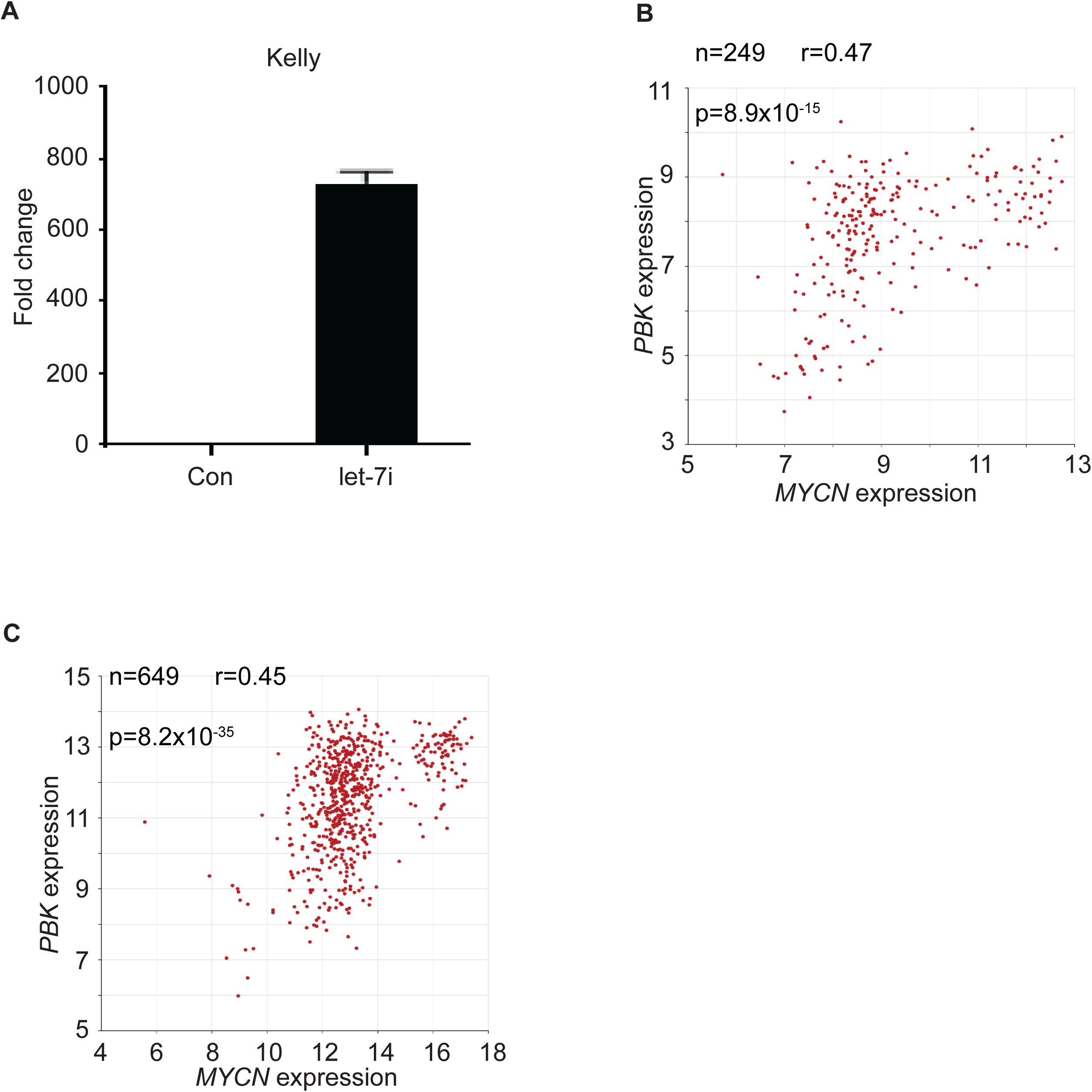
Both LIN28B/let-7 and MYCN directly regulate the expression of PBK. (A) RT-PCR demonstrates expression of let-7i in Kelly cell lines. Cell lines were infected with control lentiviruses or lentiviruses expressing let-7i. (B) Correlation between *MYCN* and *PBK* expression in 249 primary neuroblastomas. Data obtained from the TARGET dataset. (C) Correlation between *MYCN* and *PBK* expression in 649 primary neuroblastomas. Data obtained from the Kocak dataset. P values and r values listed.

**Supplementary Table S1.**
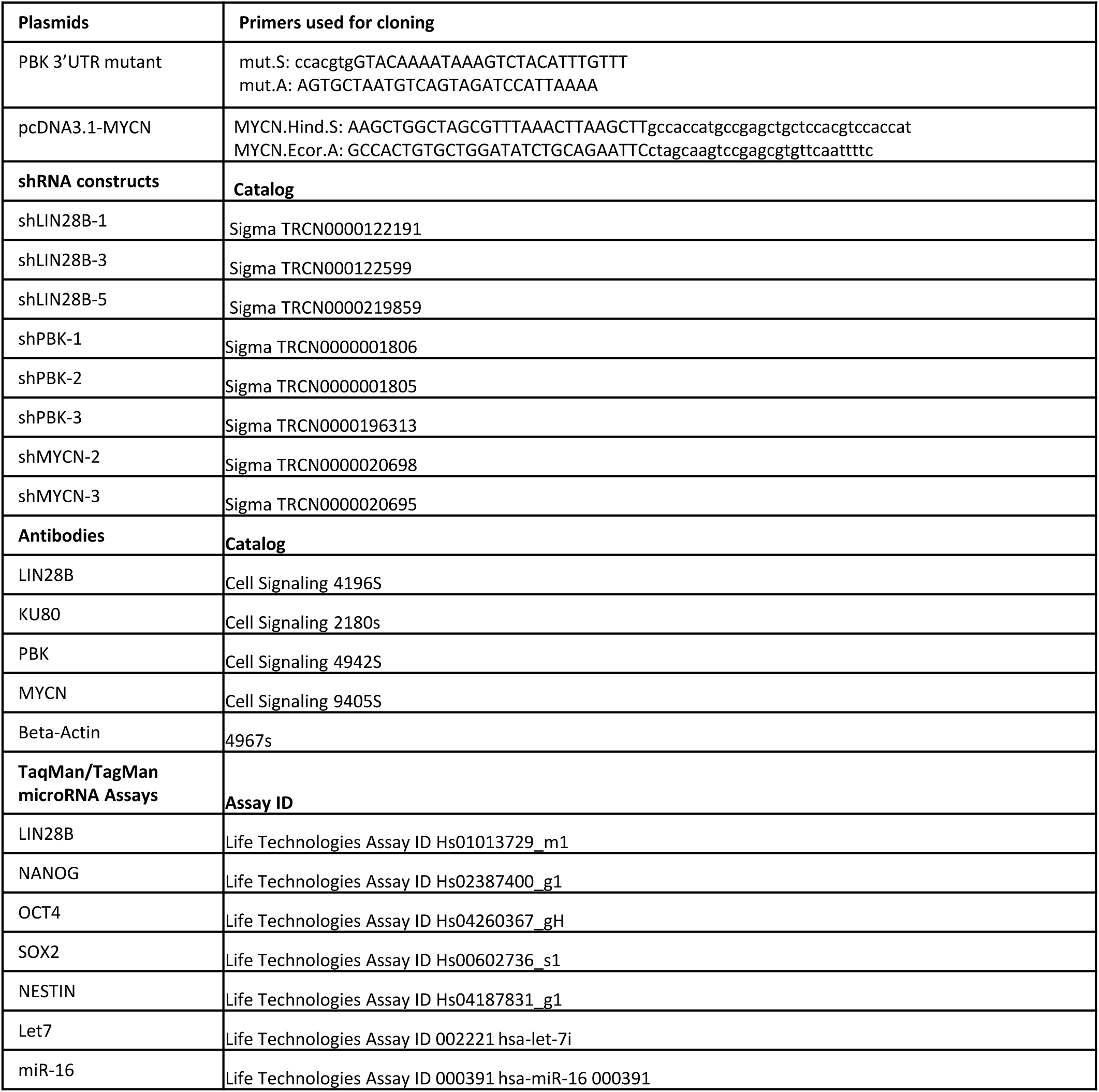
This table lists plasmids, shRNA constructs, antibodies, and TaqMan assays.

